# Feature-specific inhibitory connectivity augments the accuracy of cortical representations

**DOI:** 10.1101/2025.08.02.668307

**Authors:** Mora B. Ogando, Lamiae Abdeladim, Kevin K. Sit, Hyeyoung Shin, Savitha Sridharan, Karthika Gopakumar, Hillel Adesnik

## Abstract

To interpret complex sensory scenes, animals exploit statistical regularities to infer missing features and suppress redundant or ambiguous information. Cortical microcircuits might contribute to this cognitive goal by either completing or cancelling predictable activity, but it remains unknown whether, and how, a single circuit can implement these antagonistic computations. To address this central question, we used all-optical physiology to simulate sensory-evoked activity patterns in pyramidal cells (PCs) and somatostatin interneurons (SSTs) in the mouse’s primary visual cortex (V1). In the absence of external visual input, photostimulation of orientation-tuned PC ensembles drove either completion or cancelation of input-matching representations, depending on the number of photostimulated cells. This dual computational capacity arose from the co-existence of ‘like-to-like’ excitatory interactions between PCs, and a newly discovered ‘like-to-like’ SST-PC connectivity motif, in which SSTs are preferentially recruited by, and in turn suppress, similarly tuned PCs. Finally, we show that photoactivation of tuned SST ensembles during visual processing improved the discriminability of their preferred visual input by suppressing ambiguous activity. Thus, these complementary feature-specific connectivity motifs allow different strategies of contextual modulation to optimize inference by either completion (through PC–PC interactions) or cancelation (via PC–SST–PC loops) of predictable activity, depending on the structure of the input and the network state.

## Introduction

A central goal of neuroscience is to understand how functional interactions between neurons mediate information processing. Because the complexity of the external world vastly exceeds the computational resources of individual brains, sensory systems rely on inference to extract behaviorally relevant information. Prominent models of cortical function propose that neurons are not passive feedforward processors, but active predictors^1–3^, representing the most likely causes of incoming sensory signals. Different sensory processing frameworks propose that brain circuits use learned statistical regularities (i.e. an “internal model of the world”) to enable completion^4–9^ or cancelation^10–13^ of predictable signals (for comprehensive reviews see ^14–16)^. Each of these complementary computations can enhance inferential accuracy; for example, they can fill in missing information during occlusion or suppress redundant or ambiguous features during discrimination, respectively. However, the specific microcircuits that implement prediction cancelation or completion, and how they shape afferent inputs, remain elusive. Conflicting findings have emerged regarding how recurrent cortical networks transform orientation-tuned input patterns. While some work indicates that these circuits promote pattern completion, others imply that their primary role is to suppress tuned inputs. On one hand, co-activating large ensembles of similarly tuned PCs with two-photon (2p) holographic optogenetics generates sensory-like representations in the local network that match (i.e. “complete”) the input^4,17,18^, propagate to higher processing areas^19^ and influence decision-making^17,18^. This is consistent with prior functional and anatomical studies that have revealed enrichment in monosynaptic connections between local PCs with shared stimulus preferences^20–23^, a motif called “like-to-like” connectivity, which can enable local neurons to complete partial inputs. However, targeted photostimulation of single PCs or small PC ensembles can have the opposite effect, generating preferential suppression of co-tuned representations when paired with sensory inputs^24,25^. These latter results seemingly conflict with the like-to-like recurrent excitatory connectivity.

One possible circuit architecture that could reconcile these findings is that both excitatory–excitatory (E–E) and excitatory–inhibitory–excitatory (E–I–E) pathways follow like-to-like connectivity rules. Like-to-like E–E interactions can promote completion by amplifying weak co-tuned input patterns. In contrast, like-to-like E–I–E motifs—where the recruited inhibitory neurons suppress co-tuned excitatory neurons—could mediate suppression of redundant or ambiguous signals. Depending on which pathway is more strongly recruited, stimulation may result in either completion or suppression of similarly tuned responses, as well as intermediate outcomes. We therefore hypothesize that the network complements like-to-like excitation with feature-matched inhibitory circuits to flexibly gate these computational modes.

Computational models have shown that like-to-like inhibition can suppress redundant or ambiguous activity and improve the accuracy of sensory inferences^11,12^. In Bayesian terms, this form of prediction cancellation corresponds to *explaining away*, where the presence of one cause reduces the probability or influence of alternative causes for the same effect, effectively enhancing the differences between similar inputs. However, whether and how feature-specific inhibitory circuits actively reshape the processing of afferent inputs in cortical circuits remains a key unanswered question. While some evidence indicates that PCs preferentially innervate parvalbumin-expressing (PV) inhibitory neurons with which they share correlated visual responses^26^, PVs are weakly orientation tuned^27–30^ and this connectivity rule is not orientation-specific^26,30^. Conversely, somatostatin (SST) interneurons are nearly as orientation-tuned as PCs^31,32^, and some limited evidence suggests that they receive inputs from local co-tuned PCs^33^. However, whether SSTs reciprocally innervate PCs with which they share orientation preference will critically determine the computational role of this inhibitory feedback loop. Perhaps even more importantly, it remains unclear how these recurrent microcircuit motifs scale when more naturalistic, population-level activity patterns are engaged.

To address these key questions, we therefore sought to mechanistically understand how feature-specific excitatory-inhibitory connectivity motifs shape sensory codes via recurrent computations. For this, we combined large-scale volumetric two-photon imaging with holographically patterned optogenetics to recreate visual-like activity patterns in awake mice in the absence and presence of visual stimulation. By perturbing and monitoring the activity of excitatory and inhibitory neurons, we dissected the cell-type specific circuit mechanisms underlying recurrent effects. We reveal a new ‘like-to-like’ connectivity motif that is specifically and reciprocally shared between PC and SST neurons (but not PV neurons) that enables local feature suppression. Furthermore, enhancing this tuned SST-mediated inhibitory component during visual processing increased the accuracy of cortical representations by sharpening the population code. This work thus identifies a feature-selective PC-SST architecture that enhances orientation inference through a combination of ‘like-to-like’ excitation and ‘like-to-like-to-like’ inhibition. These findings reconcile prior contradictory results on recurrent circuits in visual computation, revealing a co-alignment of tuned excitatory and inhibitory connectivity that can support visual processing by either completing or cancelling predictable activity patterns. If the circuit motifs we revealed here are shared throughout the cortex, feature-specific recurrent inhibition from SSTs might represent a general mechanism by which the cortex flexibly augments the accuracy of neural codes.

## Results

### V1 recurrent circuits drive opponent feature-specific network effects depending on the size of the activated ensemble

Because prior studies conflict on the role of recurrent cortical circuits in V1, we hypothesized that the same cortical microcircuit could drive either feature suppression or feature completion depending on properties of the input pattern. To test this notion, we performed all-optical circuit perturbations in awake mice and aimed to locally recreate visual-like activity patterns directly in the layer 2/3 of the visual cortex in the absence of external sensory input. This allows us to determine how the local recurrent circuitry transforms specific patterns of activation, via either feature-specific completion (i.e. enhanced input-matching representations) or suppression (i.e. reduced input-matching representations) in the surrounding network.

To gain a more complete understanding of how specific parameters of photostimulation alter network outputs, we explored a large input space, probing more than 1,000 individual co-tuned ensembles across 56 sessions in 15 mice, yielding more than 70,000 neurons and more than 1,000,000 unique input-output combinations (where *input* corresponds to a stimulated ensemble and *output* is a non-targeted local PC). For this, we utilized a custom large-scale 2p multiplane calcium imaging system (see Methods) to record the visual responses of large populations of neurons (1,400 ± 62 cells per session) to gratings varying in orientation (0°, 45°, 90°, 135°). After determining each neuron’s preferred orientation online, we optogenetically reactivated ensembles systematically varying in size from 1 to 100 co-tuned PCs for each orientation using 2p holographic optogenetics (3D-SHOT^34–36)^, while mice passively viewed a gray screen (Figure 1A-D).

**Figure 1.**
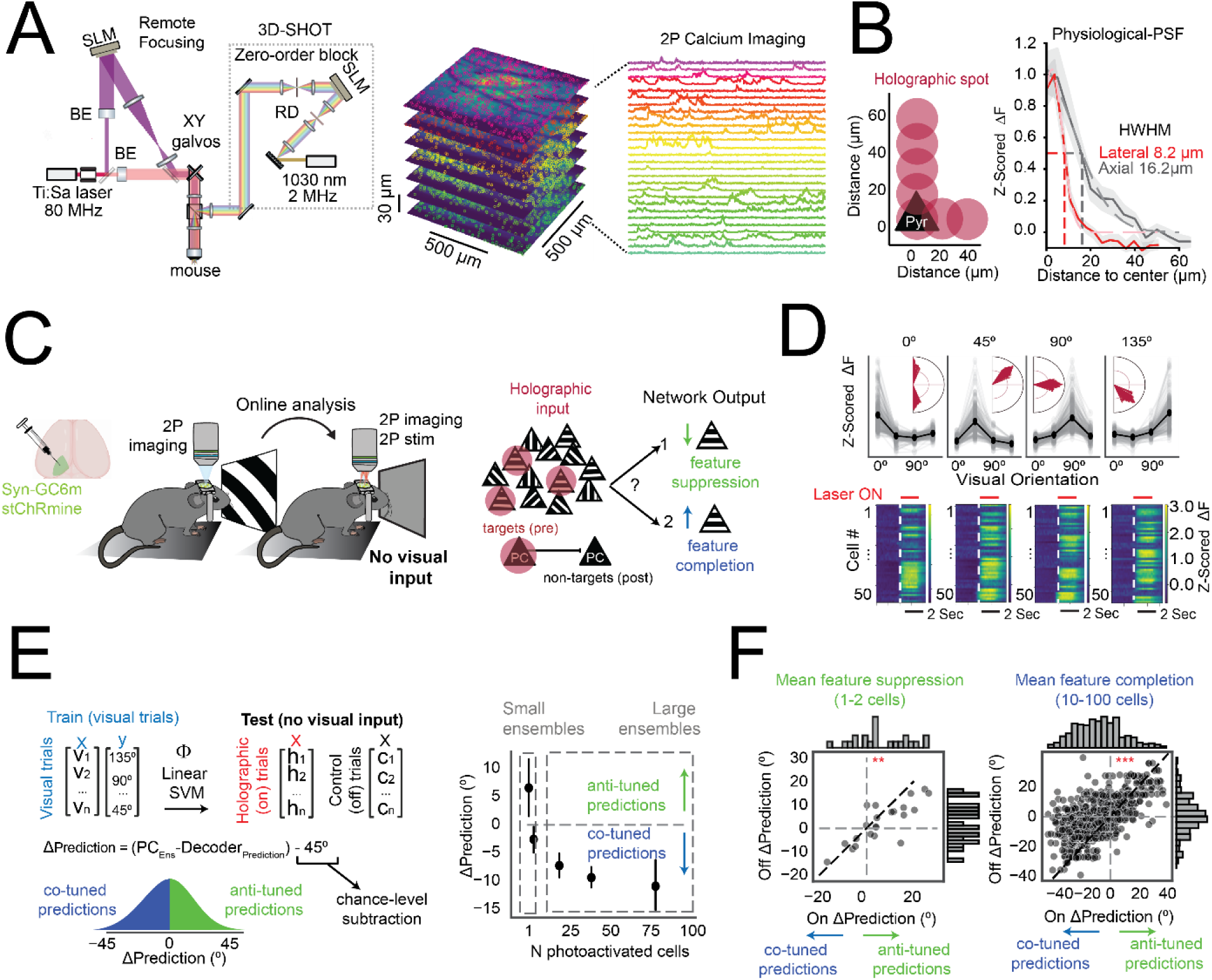
V1 recurrent circuits drive opponent feature-specific network effects depending on the size of the activated ensemble. **(A)** Schematic of the custom 3D-SHOT two-photon system, integrated with a remote focusing module for simultaneous multi-plane imaging and photostimulation (see Methods). Right: example z-stack (7 planes, 27–37 µm spacing) and representative calcium traces from neurons across cortical depth. BE=beam expander, SLM = spatial light modulator, RD= rotating diffuser. **(B)** Physiological point spread function (PSF) of holographic stimulation measured in vivo. Left: Holographic spot locations were shifted laterally and axially relative to the center of a targeted PC. Right: Average half-width at half-maximum (HWHM) across neurons: Lateral = 8.2 µm, axial = 16.2 µm; *N* = 192 neurons, 5 sessions, 3 mice). **(C)** Experimental workflow. Visual responses were first mapped using two-photon calcium imaging in awake mice expressing Syn-GCaMP6m and stChRmine. Co-tuned PC ensembles were then selectively photostimulated using 3D-SHOT in the absence of visual input. Right: Red spheres represent functionally defined, orientation-selective PCs targeted for holographic activation. We tested whether the non-targeted population was enriched in co-tuned-like (feature completion) or anti-tuned-like (feature suppression) activity. **(D)** Example visual tuning curves (top) and holographically evoked activity (bottom) from ensembles with distinct preferred orientations (0°, 45°, 90°, 135°). Each heatmap represents stimulus-aligned z-scored fluorescence of 50 targeted co-tuned PCs (rows) from an individual ensemble. **(E)** Left: Schematic of decoder assay. A linear SVM was trained to classify visual orientation based on neuronal population responses to visual stimuli (0°, 45°, 90°, 135°) and tested on non-targeted PC responses to ensemble stimulation in the absence of visual input. Middle: ΔPrediction is estimated as the difference between the predicted orientation and the input ensemble orientation. Positive values indicate feature suppression (less co-tuned classifications) while negative values indicate feature completion (more co-tuned classifications). Right: Mean ΔPrediction as a function of ensemble size. Ultra-small ensembles (1–2 cells) yielded positive ΔPrediction (preferential anti-tuned decoding; \**p* = 0.033, Wilcoxon signed-rank test, *N* = 22 ensembles, 13 sessions, 7 mice), while large ensembles (10–100 cells) produced negative ΔPrediction (preferential co-tuned decoding; \*\**p* < 0.0001, *N* = 645 ensembles, 52 sessions, 15 mice). Error bars represent 95% confidence intervals. **(F)** Comparison of decoder predictions for each ensemble during holography OFF vs. ON trials. Left: Small ensembles (1–2 cells) shift decoder classifications away from the stimulated ensemble’s preferred orientation (paired Wilcoxon signed-rank test, \**p* = 0.008, *N* = 22 ensembles). Right: large ensembles (10–100 cells) promote co-tuned classifications (\*\**p* < 0.0001, *N* = 645 ensembles). Insets: Histograms of ΔPrediction shifts for each condition.

To estimate the visual information that each photostimulated ensemble propagates to the surrounding PC population, we developed a decoder-based quantitative approach. We trained a linear Support Vector Classifier (SVC) on neuronal responses recorded during visual stimulation trials and then probed the model’s predictions with neural activity patterns generated by photostimulation in the absence of any visual stimulus. Because the probe trials lack any bottom-up sensory information, the decoder’s outputs when probed with optogenetic trials can unambiguously assess whether the targeted ensemble can propagate meaningful information to the rest of the imaged population. Importantly, this decoder was trained and probed only using activity from non-targeted neurons located outside the photoactivation zone to avoid any possible contamination from the directly photostimulated neurons (Figure 1E). We defined Δ-prediction as a measure of how much decoder classifications are biased toward or away from the preferred orientation of the photostimulated ensemble. To calculate this, we measured the angular difference between the decoder’s mean predicted orientation across trials and the preferred orientation of the photostimulated ensemble, which yields a continuous distribution between 0° for perfectly co-tuned and 90° for anti-tuned predictions. We compared this difference to the expected difference under chance-level predictions resulting in a shifted distribution between -45^0^ for co-tuned and 45^0^ for anti-tuned predictions (see Methods). In this scheme, Δ-prediction values below 0° indicate an enrichment in predictions aligned with the photostimulated ensemble (co-tuned classifications); a Δ-prediction value above 0° indicates suppression of such representations (anti-tuned classifications); while the absence of feature-specific activity propagation will elicit Δ-prediction values around 0°, indicating chance performance.

Using this strategy, we tested the hypothesis that recurrent circuits can drive either feature suppression or feature completion depending on the input configuration. Specifically, the number of active neurons differs across prior experiments^4,17–19,24^, which could lead to differences in recurrent dynamics due to cell-type specific patterns of synaptic convergence and divergence and non-linearities like action potential threshold or input/output gain^37^. Thus, we tested how the size of the photostimulated ensemble (from 1-100 neurons) accounts for different network outcomes. Despite substantial overall variability in network outcomes across ensembles (Figure S1B), activation of just 1 or 2 co-tuned neurons significantly reduced decoding predictions for their preferred orientation, indicative of feature suppression, consistent with previous findings^24^. In contrast, activation of larger ensembles (10-100 neurons) strongly enhanced decoding of the co-tuned orientation (consistent with^4,17,18,35^, Figure 1E). We validated this result with a paired analysis comparing decoder predictions for each ensemble during baseline (no photostimulation) and stimulation trials (Figure 1F). This analysis confirmed that small ensembles significantly reduced co-tuned predictions relative to each session’s baseline predictions, consistent with feature-specific suppression, while large ensembles enhanced co-tuned predictions, consistent with feature completion. Thus, these results demonstrate that cortical microcircuits in V1 can switch between driving feature-specific suppression to feature-specific completion depending on the sparsity of network activation.

### Pyramidal ensembles preferentially recruit co-tuned SST neurons

What are the functional mechanisms underlying this dual computational capacity? While feature-specific completion can be explained by the known like-to-like excitatory connections between PCs, the preferential suppression of co-tuned representations suggests the existence of tuned inhibitory connections in the same network^24,38,39^. We therefore mapped the feature-specific recruitment of both excitatory and inhibitory populations during PC ensemble stimulation. For this, we plotted the mean impact on local neurons as a function of the postsynaptic cells’ orientation preference relative to the photostimulated ensemble for small and large ensembles. We use the notation “Δθ(*x*_*Ens*_ − *Y*)” to denote the angular difference (Δθ) between the preferred orientation of an input ensemble composed of cell type X and that of an output neuron of cell type Y. For example, “Δθ(*PC*_*Ens*_ − *PV*) = 0°” refers to the responses of co-tuned (Δθ = 0°) PV neurons to PC ensembles.

We hypothesized that small input ensembles would engage PC–PC interactions differently than large ensembles, potentially leading to distinct network outcomes. We found that photostimulating large (more than 10 PCs), ensembles preferentially suppressed orthogonally tuned PCs, effectively enhancing the relative activity levels of co-tuned PCs (Δθ(*PC*_*Ens*_ − *PC*) = 0° >> Δθ(*PC*_*Ens*_ − *PC*) = 90°,***p<0.0001, Figure 2B). This is broadly consistent with prior experiments in V1 which found a relative enhancement of similar visual representations^4,17,18^, and with connectivity data^20,22,23^. In contrast, small ensembles did not differentially recruit co-tuned vs orthogonally tuned PCs (Figure 2B), suggesting that the decoder did not rely solely on global mean activity levels of co-tuned vs. anti-tuned neurons. Therefore, we reasoned that small inputs might produce subtle, yet structured modulations within a subset of decoder-informative neurons. These modulations of the network might shift the population code away from the stimulated orientation and favor anti-tuned representations. To directly test this, we estimated the population-level ‘representational evidence’ for each orientation by projecting trial-averaged population responses onto orientation-selective decoder axes (see Methods, see also^17,24^). These are dimensions of population activity that best separate visual responses to each orientation from the others. This analysis revealed that small ensembles favored anti-tuned visual representations relative to co-tuned representations. In contrast, large ensembles enhanced the co-tuned decoder representations relative to anti-tuned representations (Figure 2C, right).

**Figure 2.**
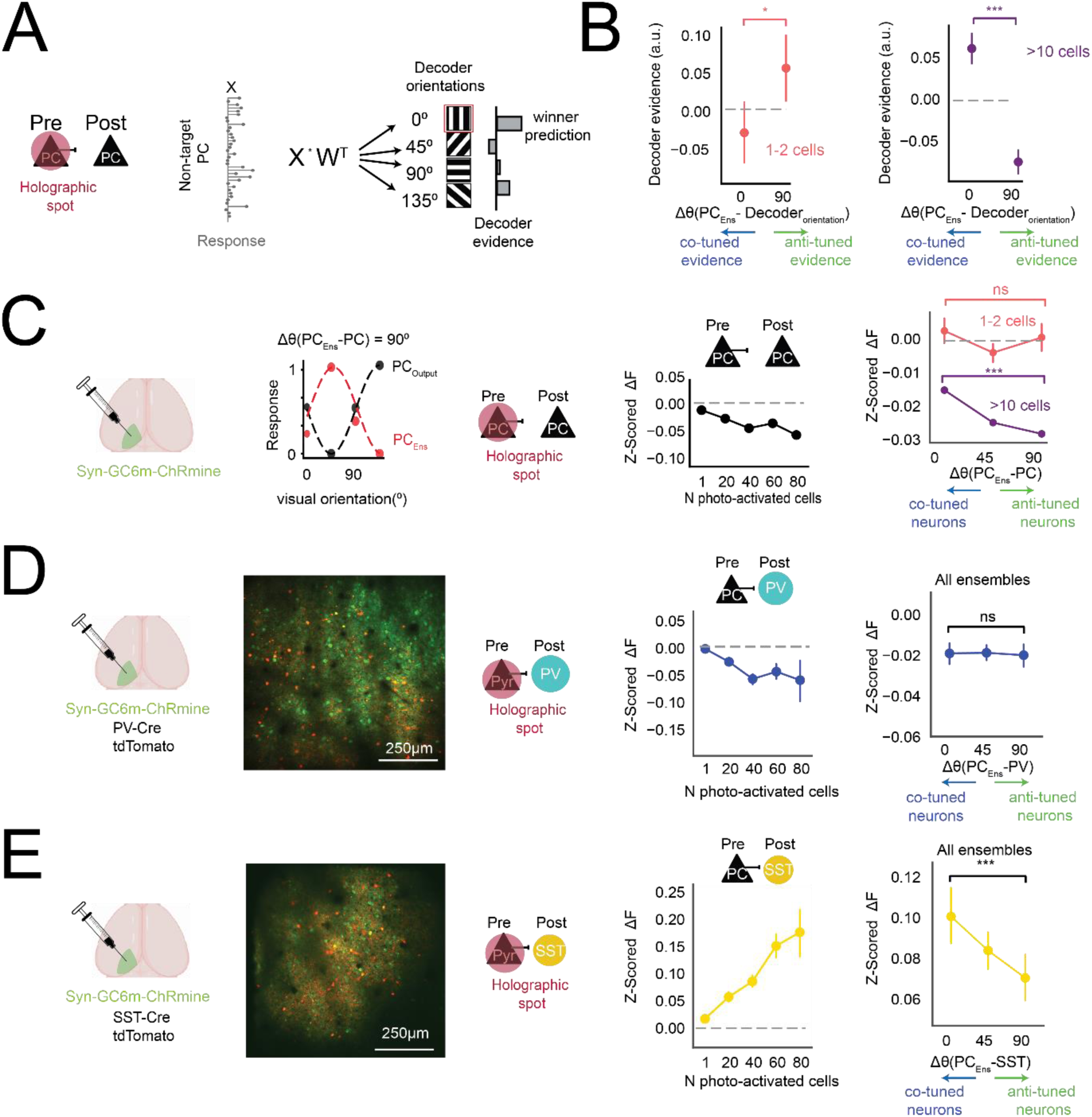
Pyramidal ensembles preferentially recruit co-tuned SST neurons. **(A)** Schematic. To quantify how holographically evoked activity resembled real visual representations, decoder evidence was computed as the average response × decoder weight across all non-targeted PCs for each decoder orientation, and plotted relative to the ensemble’s preferred orientation (Δθ(*PC*_*Ens*_ *Decoder*_*orientation*_)). **(B)** For small ensembles (1–2 cells), decoder evidence was greater for anti-tuned orientations relative to co-tuned orientations (\**p* = 0.011, paired Wilcoxon signed-rank test, *N* = 20 ensembles, 13 sessions, 7 mice), with a positive correlation between evidence and tuning offset (Δθ(*PC*_*Ens*_ − *DecOri*), Spearman *r* = 0.249, \**p* = 0.026). In contrast, large ensembles (≥10 cells) enhanced decoder evidence for co-tuned orientations relative to anti-tuned orientations (\*\**p* < 0.0001, paired Wilcoxon signed-rank test, *N* = 645 ensembles, 52 sessions, 15 mice), consistent with feature-specific pattern completion. Error bars represent 95% confidence intervals. **(C)** Left: Experimental design. Mice were injected with a pan-neuronal virus expressing GCaMP6m and st-ChRmine. Example tuning curves illustrate an angular difference between a PC and a stimulated ensemble. Middle: Mean population responses in non-targeted PCs across different ensemble sizes. Photostimulation produced net suppression in surrounding PCs, which increased as a function of the number of targeted cells (black). Right: Responses of non-targeted PCs as a function of the angular difference between each PC’s preferred orientation and that of the stimulated ensemble (Δθ(*PC*_*Ens*_ − *PC*)). For large ensembles (purple, ≥10 cells), anti-tuned responses were significantly more suppressed than co-tuned responses (Δθ(*PC*_*Ens*_ − *PC*) 0° vs. 90°; ***p < 0.001, permutation test), with a significant negative correlation with Δθ (Spearman *r* = –0.0318, *p* = 1.07 × 10⁻⁹¹; *N* 406,522 cell–ensemble comparisons across Δθ). In contrast, small ensembles (pink, 1–2 cells) showed no evidence of tuning-dependent suppression: Neither correlation nor pairwise tests between co-tuned and anti-tuned groups were significant (Spearman *r* = –0.0145, *p* = 0.13; permutation test Δθ = 0° vs. 90°: *p* = 0.45; *N* > 2523). Error bars represent 95% confidence intervals. **(D)** PV-Cre;tdTomato mice were used to label PV interneurons (red), while GCaMP6m and st-ChRmine were expressed pan-neuronally, enabling simultaneous optical stimulation of PC ensembles and calcium imaging of PV and PC responses. Left: Representative field of view showing co-expression of GCaMP6m and tdTomato. Center: PV interneurons exhibited net suppression in response to nearby pyramidal cell ensemble stimulation. Right: PV responses did not show orientation-specific modulation. There was no significant correlation between PV response amplitude and tuning offset (Δθ(*PC*_*Ens*_ − *PV*), Spearman *r* = –0.0001, *p* = 0.994; total *N* = 5948 cell–ensemble comparisons). Permutation test across Δθ bins (*p* = 0.6891), with group sizes of *N* > 1454 (Δθ = 0°). Error bars represent 95% confidence intervals. **(E)** SST-Cre;tdTomato mice were used to label SST interneurons (red), and pan-neuronal expression of GCaMP6m and st-ChRmine enabled simultaneous photostimulation of PC ensembles and calcium imaging of SST and PC responses. Left: Representative field of view showing co-expression of GCaMP6m and tdTomato. Center: SST interneurons exhibited robust activation in response to PC ensemble stimulation, with response amplitude increasing as a function of ensemble size. Right: SST responses were feature-specific, showing stronger activation when their orientation preference was aligned with that of the stimulated ensemble. This tuning-dependent modulation was supported by a significant negative correlation between SST response and tuning offset (Δθ(*PC*_*Ens*_ −*SST*), Spearman r = –0.035, \*\**p* = 0.0052; *N* = 6346 cell-ensemble combinations), and by a permutation test showing significant differences in SST responses across Δθ bins (\*\*\**p* = 0.0002). Error bars represent 95% confidence intervals.

In all cases, we observed mean suppression of nearby PCs (Figure S2, see also^24,39^) across ensemble sizes, supporting the idea that local recurrent inhibition (rather than excitation) is the dominant influence PCs exert on surrounding PCs. Previous work has shown that trains of action potentials in individual PCs can robustly evoke responses in SST cells^33,40,41^, but less so in PVs^41^. Moreover, unlike PVs, SST interneurons are nearly as orientation-selective as PCs^31,32,42^ (Figure S2 F-H). However, whether the activation of co-tuned PCs at the ensemble level differentially recruits PV vs. SST subnetworks, as well as their feature-specific connectivity rules remain completely unexplored. To address this, we transgenically labeled PVs or SSTs, and stimulated ensembles of PCs while monitoring activity in these GABAergic subtypes (see Methods, Figure 2 C,D).

Using this approach, we first quantified the recruitment of the two subpopulations as a function of the number of photostimulated PCs. Surprisingly, despite abundant PC-PV inputs in this network (revealed by physiological and anatomical studies, see^30,43^), photostimulation of local PC ensembles suppressed PV activity (Figure 2C, left panel), mirroring the mean suppression of the local PC population (Figure 2A). In striking contrast, PC-ensemble photostimulation strongly recruited SSTs, with SST activation scaling proportionally with the number of stimulated PCs (Figure 2D, left panel). This strongly suggests that SSTs are the dominant drivers of the recurrent suppression seen above across all ensemble sizes, consistent with one photon stimulation of L2/3 PCs^44^.

Can this inhibitory population drive feature-specific suppression? We found that SST recruitment strongly depended on their shared orientation preference with the input ensemble, consistent with recent findings at the single-cell level^33^ (responses for Δθ(*PC*_*Ens*_ − *SST*) = 0° vs. responses for Δθ(*PC*_*Ens*_ − *SST*) = 90°,***p<0.0001, permutation test, Figure 2D, right panel). In contrast, the mean suppressive effect on PVs did not depend on their shared orientation preference with the photostimulated PC ensemble, consistent with reports using paired intracellular recordings^26^ (Figure 2C, right panel). To our knowledge this is the first functional demonstration of the cell-type specific recruitment of tuned inhibition by cortical PC ensembles. This result suggests that the orientation preference of SST neurons might arise from their local PC inputs in this layer, and, through recurrent feedback inhibition, SSTs might exert orientation-specific suppression of the network. Notably, while PCs preferentially recruited co-tuned SSTs, all SSTs strongly tracked the number of photostimulated neurons, with large ensembles activating both co-tuned and anti-tuned SSTs (Fig. S2E). This suggests that SST neurons simultaneously integrate both the average population activity and the feature alignment of PC inputs.

### Orientation-selective SST-to-PC inhibition mediates local feature suppression

Although the functional circuit analysis above shows that co-tuned ensembles of PCs activate co-tuned SSTs, it remains unclear how feature-selective inhibition from SSTs, in turn, shapes the divergent outcomes of network activation: either feature suppression or completion. To answer this question, it is necessary to determine the structure of the connectivity from SST back onto PCs. If SSTs preferentially suppress co-tuned PCs, small PC ensembles may drive feature suppression via a ‘like-to-like-to-like’ di-synaptic loop. Conversely, if SSTs preferentially suppress orthogonal PCs, large PC ensembles could enhance feature completion via a ‘like-to-like-to-unlike’ motif (Figure 3A, schematic). Additionally, in any scenario, the relative balance of tuned inhibition, untuned inhibition, and like-to-like excitation between PCs will be critical.

**Figure 3.**
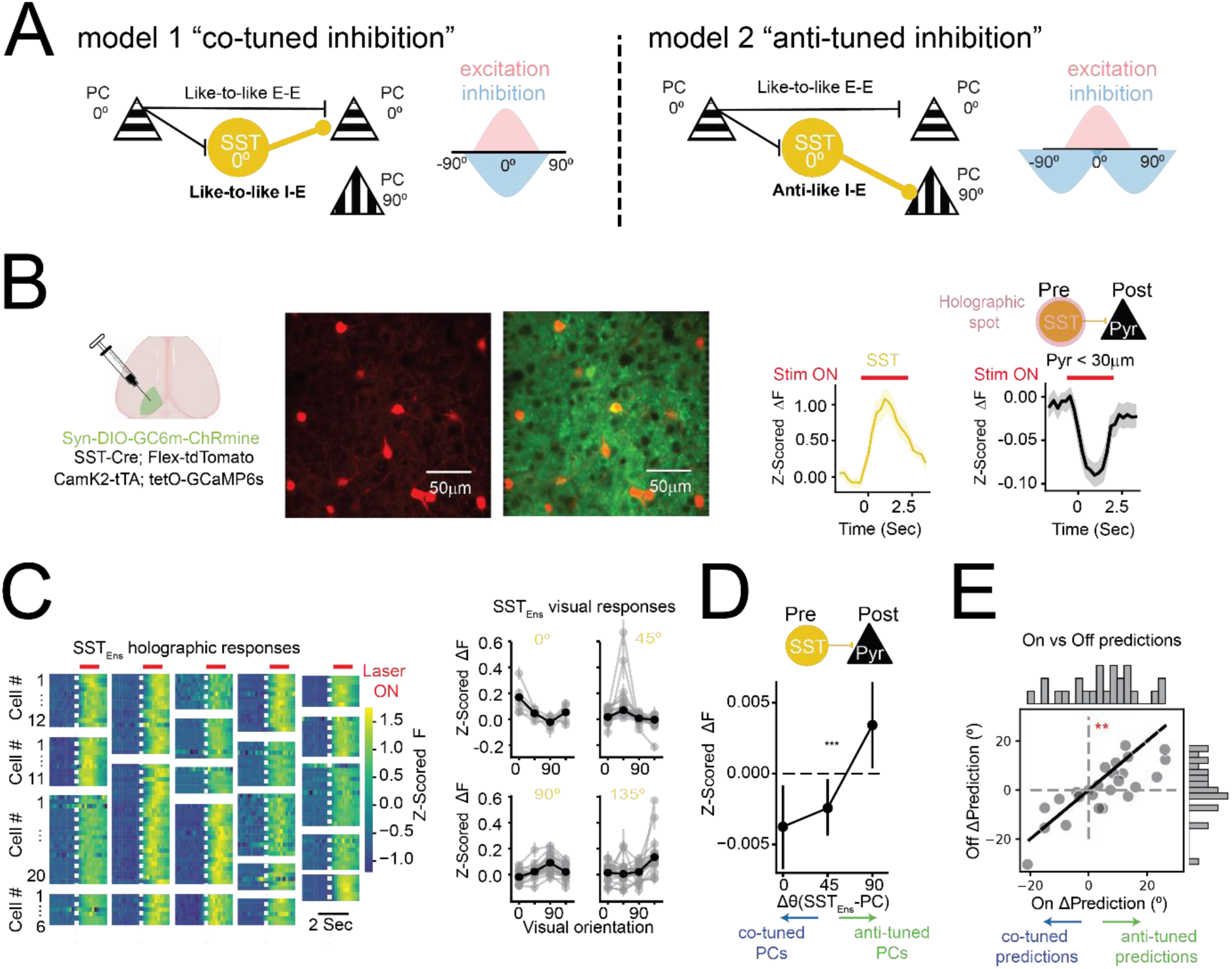
Co-tuned SST-to-PC inhibition mediates *de novo* feature suppression. **(A)** Schematic of two alternative inhibitory microcircuit motifs. Left: A ‘*co-tuned’* inhibitory motif where SST interneurons preferentially inhibit PCs with similar orientation tuning, enforcing within-feature competition. Right: An ‘*anti-tuned’* inhibitory motif where SST neurons suppress PCs tuned to orthogonal features, thereby mediating cross-feature competition. **(B)** Left: Experimental strategy using SST-Cre; Flex-tdTomato; CamK2-tTA; tetO-GCaMP6s mice. SST neurons were selectively transduced with Syn-DIO-GCaMP6m-ChRmine, allowing simultaneous calcium imaging and two-photon holographic stimulation of SST ensembles. PCs expressed GCaMP6s via the tetO system. Example session showing mean SST responses (yellow, targeted neurons) and nearby PC responses (black) to SST activation. Strong suppression in PCs near the targeted SST somas confirms the cell-type specificity of the approach. **(C)** Left: Heatmaps showing SST responses following ensemble photostimulation. Each panel represents a single ensemble with rows corresponding to individual SST neurons. Right: Orientation tuning of the same SST ensembles across four visual conditions (0°, 45°, 90°, 135°). **(E)** SST activation significantly suppressed co-tuned PCs more than orthogonally tuned PCs. This tuning-dependent modulation was supported by both a significant correlation between Δθ(*SST*_*Ens*_ − *PC*) and PC responses (Spearman r = 0.019, \*\**p* = 0.0047; *N* = 21502 cell-ensemble combinations), and by a permutation test showing significant differences in PC responses across Δθ bins (Δθ 0° vs. 90°; **p = 0.0021 N>5254 cell-ensemble combinations). Error bars represent 95% confidence intervals. **(E)** Decoder trained on visually evoked PC responses was tested on trials with and without SST ensemble photostimulation (no visual input). Each point represents the mean ΔPrediction on a single ensemble, defined as the trial-averaged angular distance between the predicted orientation and the SST ensemble’s preferred orientation, centered around chance level. Decoder predictions were significantly biased away from the stimulated SST ensemble’s preferred orientation during photostimulation (**p = 0.0076, Paired *t*-test; Shapiro-Wilk normality *p* > 0.3). Insets: Histograms of ΔPrediction shifts for each condition.

Testing these alternative circuit motifs requires precise photostimulation of co-orientation tuned ensembles composed exclusively of SST neurons, while recording the activity of nearby PCs. This required a genetically and optically precise strategy: We expressed an opsin selectively in SST neurons while labeling both SSTs and PCs with GCaMP6 (Figure 3B, also see Methods). Moreover, we leveraged our custom multiplane all-optical system with simultaneous 3D stimulation and readout across 180µm of cortical depth (see Methods). Using these tools, we identified and stimulated ensembles of approximately 10 co-tuned SST neurons per experiment (10.8 ± 1.1 SST cells, range: 6-27), drawn from an average of 45 individually resolved SSTs co-expressing GCaMP6 and opsin (Figure 3C).

Photostimulating co-tuned SST ensembles preferentially suppressed the activity of co-tuned PCs (responses for Δθ(*SST*_*Ens*_ − *PC*) = 0° vs. responses for Δθ(*SST*_*Ens*_ − *PC*) = 90°, ***p<0.0001, Figure 3D). These results causally establish feature-specific, like-to-like SST-to-PC inhibition and rule out anti-like inhibition. To our knowledge this is the first direct functional demonstration of co-tuned inhibitory→excitatory connectivity in neocortical circuits. Is this suppression sufficient to bias decoder classifications against co-tuned orientations in the absence of visual input? To test this, we trained a decoder on visually evoked PC activity, and then probed it with the PC responses during SST ensemble photostimulation (as above, in the absence of any visual stimulus). We found that SST ensemble activation reduced the probability of co-tuned classifications (Figure 3E), demonstrating that co-tuned SST activity alone is sufficient to suppress co-tuned representations in the local network in the absence of any excitatory drive.

Because co-tuned SSTs are preferentially activated by PC ensembles, and they in turn suppress similarly tuned PCs, the di-synaptic PC-SST-PC connectivity results in ‘like-to-like-to-like’ interactions. This establishes a mechanism by which small PC ensemble stimulation can induce feature suppression, if the relative recruitment of this tuned inhibitory pathway dominates over direct PC-PC interactions.

### Co-tuned inhibition can sharpen tuned inputs in a recurrent model

The preceding experiments map the functional connectivity from PCs to SSTs and PVs, and back from SSTs to PCs, and demonstrate how photostimulation of co-tuned PC or SST ensembles can influence visual-like representations in the absence of any real visual stimulus. However, it remains unclear how this feature-specific inhibition might shape the processing of incoming visual inputs. To understand the functional implications of this like-to-like SST connectivity motif, we built a recurrent network model incorporating both tuned excitation and tuned inhibition. We aimed to recapitulate the divergent network results observed in Figure 1 and then used this model to investigate how the network transforms feedforward inputs. We modeled a population composed solely of orientation-tuned PCs, incorporating both excitatory and inhibitory influences as net postsynaptic effects, with inhibition implemented as negative interactions between PCs rather than through an explicit inhibitory population (Figure 4A, see Methods). First, we searched for the conditions in which the model produced opposite feature-specific behaviors as the ones observed under PC stimulation. For this, we scaled the excitatory and inhibitory connection strengths (Figure 4B, middle; see Methods) in order to slow down the initial growth of the excitatory drive relative to the inhibitory drive as a function of the number of active cells, which is consistent with the stronger and denser connectivity between PCs and SSTs as well as the higher excitability of SSTs^41,45^. Using this strategy, we observed that in sparse stimulation conditions, where only 1-2 PCs were activated, tuned inhibition dominated and suppressed co-tuned neurons (Figure 4B). In contrast, when denser patterns of co-tuned PCs were stimulated (25 PCs), excitatory drive outcompeted inhibition, amplifying co-tuned responses, and recapitulating our experimental phenomenology under PC ensemble stimulation.

**Figure 4.**
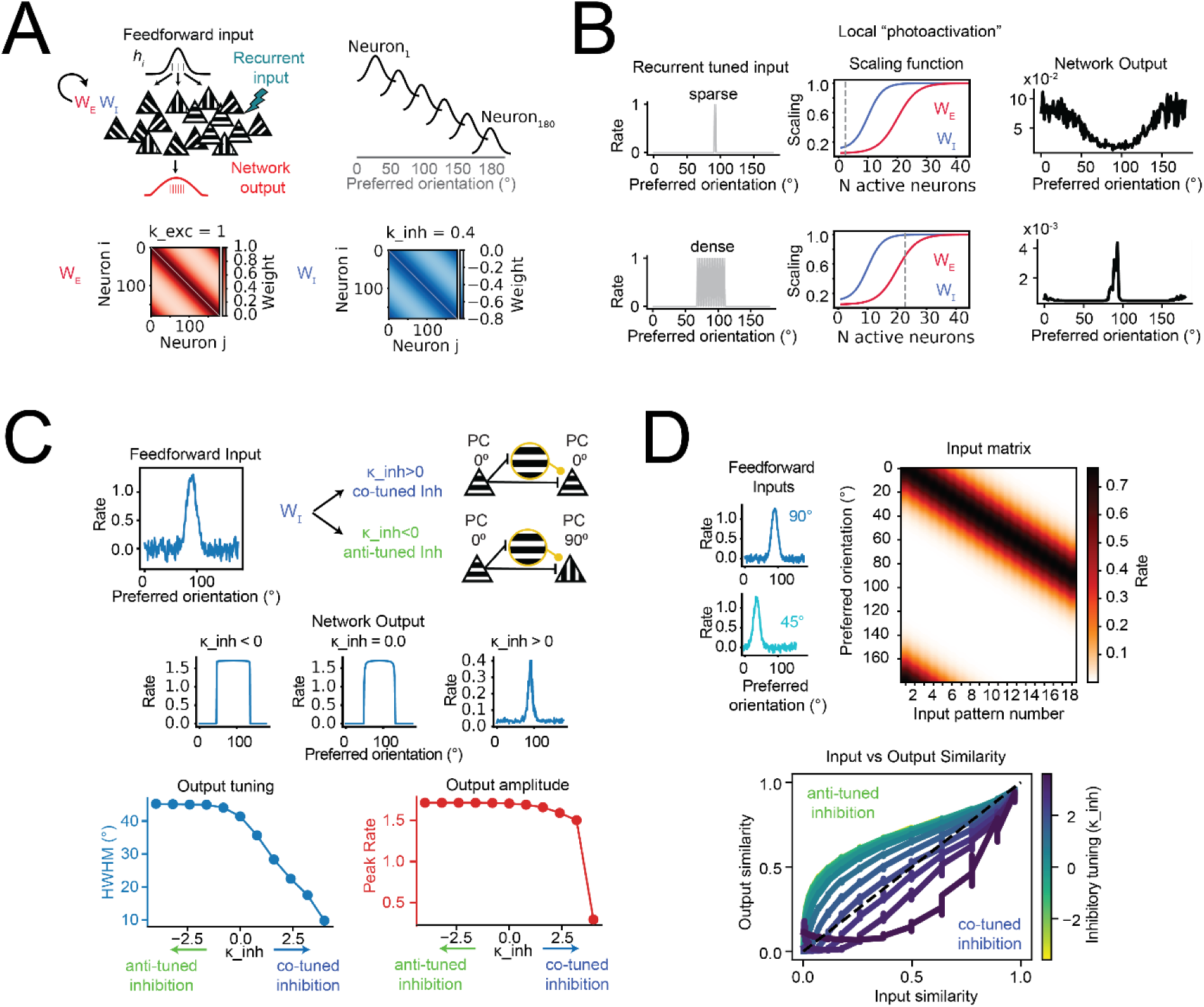
Modeling the functional implications of tuned inhibition. **(A)** Schematic of the recurrent network model. Neurons receive feedforward input and recurrent excitation and inhibition, with weights (WE and WI) shaped by the similarity in preferred orientation. Bottom: Example excitatory (left) and inhibitory (right) connectivity matrices. **(B)** Simulated local “photostimulation” of subsets of PCs, revealing differences in network response under sparse (top) versus dense (bottom) conditions. Left: input PCs with orientation preferences centered at 90° were clamped at high firing rates during the recurrent loop. Middle: scaling functions controlling excitation and inhibition as a function of network activity. Dotted line indicates the number of activated PCs for each condition (sparse vs dense). Right: resulting network output rate as a function of preferred orientation of non-targeted neurons. **(C)** Top: Feedforward inputs were simulated for models with a range of inhibitory tuning (k_inh) values. Positive k_inh denote co-tuned inhibitory connectivity, while negative k_inh indicate anti-tuned inhibitory connectivity. Middle: Example network outputs for different k_inh values. Bottom-Left: Increasing inhibitory co-tuning sharpens output tuning, measured as half-width at half-maximum of the output population tuning curve, HWHM. Bottom-Right: Highly co-tuned inhibitory connections lead to reduced peak firing rates. **(D)** Top: Feedforward inputs ranging in peak orientations from 0° to 90° and spaced by 5° were simulated for models with a range of inhibitory tuning (k_inh) values. Right: The input matrix shows the simulated patterns sorted by their peak orientation value. Bottom: Cosine similarity between input and output representations as a function of inhibitory tuning. Similar input patterns were decorrelated by networks with co-tuned inhibition.

Previous theoretical and experimental studies have shown that inhibition co-aligned with excitation can enhance the selectivity of neuronal responses by biasing the excitatory-inhibitory ratio towards excitation for the preferred orientation as compared to flanking orientations^46–49^. To test this in our model, we simulated tuned feedforward visual inputs while systematically varying the specificity of the inhibitory connectivity. In networks with anti-tuned inhibition, co-tuned excitatory neurons rapidly recruited adjacently tuned PCs (i.e., neurons preferring flanking but non-matching orientations), resulting in strong but broad population responses, quantified as half-width-half-max of the output tuning curve (Figure 4C). In contrast, co-tuned inhibition prevented this broadening in feature space, because only the neurons strongly aligned with the input peak received sufficient afferent excitation to overcome the co-tuned inhibition (Figure S3B). As a result, co-tuned inhibitory connectivity preferentially reduced “flanking activity” and sharpened the population response by restricting activation to the most stimulus-aligned subset of neurons, effectively enhancing selectivity (Figure S3B). However, when tuned inhibition was too sharp, the peak response decreased substantially (Figure 4C), which could lead to a prohibitively low signal-to-noise regime.

We reasoned that this sharper input representation might help decorrelate similar input vectors into more distinct output patterns, which can improve discrimination in downstream regions^50,51^. To quantify this, we calculated the cosine similarity between pairs of inputs and pairs of output activity patterns. We used a set of tuned inputs with peak orientations ranging from 0° to 90° to directly quantify how networks transformed inputs with different degrees of similarity (from very similar to very dissimilar). This analysis revealed that networks with co-tuned inhibition were more effective at decorrelating similar inputs into more separable output vectors (Figure 4D, bottom). These results suggest that feature-selective inhibition aligned to the stimulus could sharpen population tuning and reduce the overlap between similar representations, thereby enhancing discriminability.

### Tuned recurrent inhibition improves feature discrimination

To experimentally test these computational predictions, we optogenetically stimulated co-tuned SST ensembles varying in their orientation preference while presenting visual gratings of multiple orientations. This approach allowed us to assess whether the network effects of SST activation depend on the alignment between the SST ensemble’s preferred orientation, the visual stimulus, and the postsynaptic PC’s orientation preference. This results in three distinct orientation relationships that must be clearly distinguished. First, as before, the angular difference (Δθ) between the preferred orientation of an input ensemble (composed of SST interneurons) and that of an output PC: Δθ(*SST*_*Ens*_ − *PC*). Second, the angular difference (Δθ) between the visual stimulus and the preferred orientation of the output PC: Δθ(VisStim − PC). This parameter defines a population tuning curve, where neurons responding maximally (to their preferred stimulus) are considered “on-peak,” while others contribute to the “off-peak” response. Finally, the angular difference (Δθ) between the orientation of the visual stimulus and the preferred orientation of an input ensemble (composed of SST interneurons): Δθ(VisStim − *SST*_*Ens*_). We use this parameter to distinguish trials where SST photostimulation reinforces the natural activation of co-tuned inhibitory neurons (“SST-aligned”) from those where it recruits misaligned ensembles not normally engaged by the visual input (“SST-misaligned”). Moreover, to evaluate how SST perturbations modulate the shape of tuning curves across PCs with varying maximum response amplitudes, we normalized each PC’s activity to its response to the preferred visual stimulus (peak response) in visual-only trials. With this normalization, a peak value of 1 indicates that the PC responds equally in the presence (SST-ON) and absence (SST-OFF) of SST stimulation.

First, we quantified the mean responses of all PCs as a function of the Δθ(*SST*_*Ens*_ − *PC*), independent of the visual input. In agreement with our previous results, we found that SSTs suppressed co-tuned PCs more than anti-tuned PCs on average (*p<0.05, for Δθ(*SST*_*Ens*_ − *PC*) 0° vs. 90° across all visual stimuli, permutation test). However, when considering the alignment between each PC’s preferred orientation and the visual input (Δθ(VisStim − PC)), we observed that this differential suppression in PC populations occurred only when PCs responded to sub-optimal stimuli, not matching their preferred visual input (Figure 5C, left). At their peak (when Δθ(VisStim − PC) = 0°), both co-tuned PCs (Δθ(*SST*_*Ens*_ − *PC*) = 0°) and anti-tuned PCs (Δθ(*SST*_*Ens*_ − *PC*) = 90°) showed moderately reduced responses compared to visual-only trials, with co-tuned PCs being slightly less suppressed (responses below 1 at the peak, Figure 5C left). This interaction between the optogenetic perturbation and the visual input is consistent with recent findings showing that both the strength of the visual input as well as the brain state can shift the net effect of SST recruitment on the local network, possibly due to SST-dependent network stabilization^52^.

**Figure 5.**
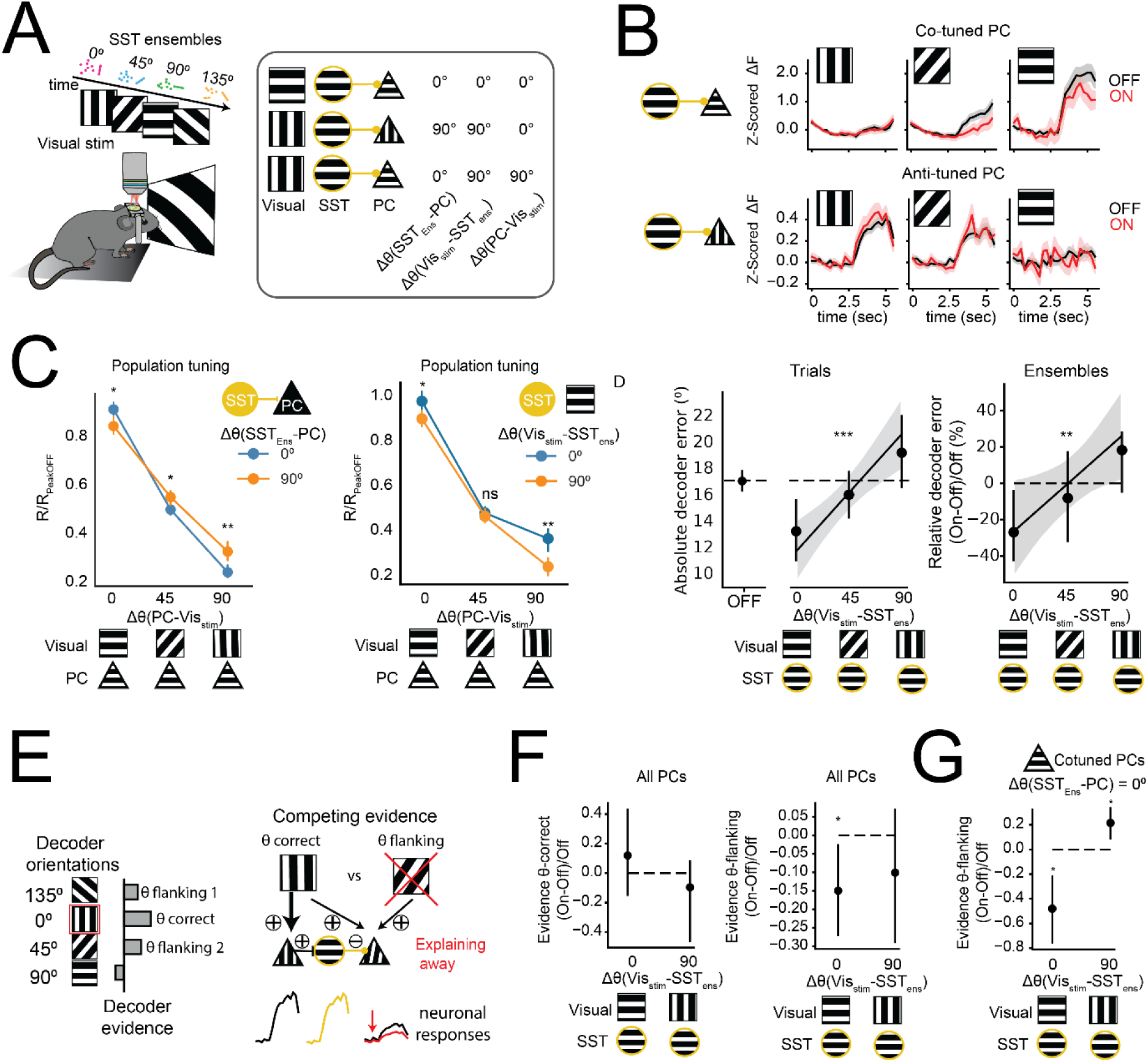
Tuned recurrent inhibition improves feature discrimination. **(A)** Schematic illustrating the experimental strategy: visual stimuli were paired with optogenetic activation of a tuned SST ensemble. Right, scheme showing the three distinct orientation relationships analyzed across the visual stimulus, the stimulated ensemble and the local PC: Δθ(*SST*_*Ens*_ − *PC*), Δθ(*Vis*_*Stim*_ − PC), Δθ(*Vis*_*Stim*_ − *SST*_*Ens*_). Different combinations of these parameters are shown in three illustrative examples. **(B)** Example PC responses during SST-OFF (black) and SST-ON (red) trials for co-tuned (top) and anti-tuned (bottom) PCs, show stimulus-specific suppression during SST activation for the co-tuned neuron. **(C)** Left: Normalized population tuning curves (median ± 95CI) of PCs grouped by their relative visual preference (Δθ(*PC* − *Vis*_*Stim*_), 0°, 45°, or 90°). Colors indicate tuning alignment between the stimulated SST ensemble and the recorded PC: Δθ(*SST*_*Ens*_ − *PC*) = 0° or 90° correspond to blue and orange respectively. SST activation depends on both visual and ensemble alignment. At Δθ(*PC* − *Vis*_*Stim*_) >0, co-tuned PCs (Δθ(*SST*_*Ens*_ − *PC*)= 0°) showed significantly lower responses compared to anti-tuned PCs (Δθ(*SST*_*Ens*_ − *PC*)= 90°; permutation test: **p = 0.006 and ***p = 0.0002, for Δθ(*PC* − *Vis*_*Stim*_) = 45° and Δθ(*PC* − *Vis*_*Stim*_) = 90° respectively). In contrast, at Δθ(*PC* − *Vis*_*Stim*_) = 0°, suppression was weaker in co-tuned compared to anti-tuned PCs (**p = 0.009). Right: Normalized population tuning curves (median ± 95CI) of PCs grouped by their relative visual preference (Δθ(*PC* − *Vis*_*Stim*_), 0°, 45°, or 90°). Colors indicate tuning alignment between the stimulated SST ensemble and the visual input: Δθ(*Vis*_*Stim*_ − *SST*_*Ens*_) = 0° or 90° correspond to blue and orange respectively. Permutation test: *p = 0.002, p>0.05 and ***p < 0.0001, for Δθ(*PC* − *Vis*_*Stim*_) = 0°, 45° and 90° respectively. **(D)** SST stimulation reduces decoder errors in visually aligned inputs. Left: Mean decoder error (in degrees) for SST-OFF trials. Middle: Mean decoder error for interleaved SST-ON trials, grouped by alignment between the SST ensemble and the visual input (Δθ(VisStim − *SST*_*Ens*_)). Asterisks indicate a significant correlation between Δθ(*Vis*_*Stim*_ − *SST*_*Ens*_) and decoder error (Spearman r = 0.09, ***p = 0.0001, N=1987 trials, from 6 mice). Right: Percentage of change in decoder errors (median ± 95CI) relative to SST-OFF trials. Stimulus-aligned SST activation Δθ(*Vis*_*Stim*_ − *SST*_*Ens*_)= 0° reduces decoder errors, indicating enhanced discrimination (*p = 0.041, permutation test, N = 23 ensembles, 9 sessions, 6 mice). Asterisks indicate a significant correlation between Δθ(*Vis*_*Stim*_ −*SST*_*Ens*_) and decoder error (Spearman r = 0.27, **p = 0.009). **(E)** Schematic representing decoder-based evidence analysis quantifying “explaining away” (adapted from^12^). Explaining away is computed as the reduction in evidence for the orientations flanking the stimulated visual input, in SST-ON trials compared to SST-OFF trials. **(F)** Left: Evidence for the correct orientation during SST-ON does not change for any visual condition, relative to SST-OFF trials. Right: SST activation selectively suppresses evidence for flanking orientations in aligned visual inputs (Δθ(*Vis*_*Stim*_ − *SST*_*Ens*_) = 0°: *p=0.032, permutation test). **(G)** SST photoactivation reduces the relative decoder evidence for flanking orientations carried by co-tuned PCs (Δθ(*SST*_*Ens*_ − *PC*)= 0°) and in aligned visual trials (Δθ(*Vis*_*Stim*_ − *SST*_*Ens*_) = 0°, *p=0.025), but not in misaligned trials (Δθ(*Vis*_*Stim*_ − *SST*_*Ens*_) = 90°, *p=0.025 permutation test).

Can this population-specific and visual-specific modulation change the discriminability of visual representations? If SSTs inhibit co-tuned neurons only when the stimulus is misaligned with their preference (during off-peak responses), then decoding performance should improve when SSTs are *not* aligned with the stimulus, since this implies a reduction in inappropriate responses (Figure 5C, right). However, our modeling results suggest that anti-tuned inhibition is less effective than co-tuned inhibition at decorrelating *similar* orientation tuned inputs. We reasoned that if discrimination errors typically occur between similar orientations, boosting the natural recruitment of co-tuned SST ensembles might contribute to the separation of these population responses, improving performance.

To test this, we trained a decoder on visual-only trials and compared decoder performance on interleaved trials with and without SST activation. As above, this decoder was trained and tested using non-photostimulated neurons. We quantified each SST ensemble’s alignment to the stimulus using the absolute orientation difference between the ensemble’s preferred tuning and that of the visual stimulus (Δθ(VisStim − *SST*_*Ens*_)) as explained above. Importantly, this metric depends exclusively on the SSTs and the visual input and does not incorporate the tuning of the PCs. Consistent with our modeling results, we observed that average decoder error (relative to interleaved visual-only trials) was significantly reduced in SST-aligned trials by ∼25% (Figure 5D right, 27% ± 8% for Δθ(VisStim − *SST*_*Ens*_) < 45°, *p = 0.04, permutation test), but not in SST-misaligned trials (Δθ(VisStim − *SST*_*Ens*_) > 45°). To confirm this tuning-dependent benefit, we found a significant positive correlation between Δθ(VisStim − *SST*_*Ens*_) and decoder error change (Spearman R = 0.27, **p =0.009). Finally, restricting the analysis to the same SST ensembles paired with opposite visual stimuli, we found that decoder errors were significantly lower when the visual stimulus was aligned with the ensemble’s tuning (*p= 0.02, Wilcoxon signed-rank test). These results demonstrate that SST-mediated inhibition enhances visual orientation coding specifically when the SST ensemble’s orientation preference matches the visual stimulus.

What is the mechanism underlying this improved linear discriminability? One possibility is that SST neurons influence population responses in a specific subset of PCs that the decoder is reading from (“decoder-informative” PCs). Alternatively, rather than modifying mean responses, SST-activation might differentially remove activity shared across multiple inputs, enhancing the distinct components that are unique to the presented stimulus, while maintaining overall activity levels. To probe these scenarios, we first plotted the mean population tuning curves using solely decoder-informative PCs (neurons with non-zero weights after L1 regularization, see Methods) in SST-aligned vs SST-misaligned trials. Interestingly, this subgroup of PCs responded equally in SST-aligned compared to misaligned trials across visual conditions (Figure S4A). To explore whether these population responses were more distinct for different visual inputs, we quantified the cosine similarity between flanking (45° apart) visual orientations, using decoder-informative PCs. We observed a significant reduction in similarity on SST-aligned trials compared to SST-misaligned trials (Figure S4B), indicating that SST stimulation can reshape population activity in an input-dependent manner, without altering overall activity levels.

This reduction in similarity across similar visual inputs is consistent with the principle of “explaining away” in which the network actively downregulates evidence for alternative interpretations of ambiguous input, increasing the relative weight of the most likely cause (Figure 5E). Several computational models have employed within-feature competition to improve inference through explaining away^11,12,24^, but a causal demonstration of this computation in neocortical circuits has not yet been achieved. We quantified how SST activation altered the decoder ‘representational evidence’ for visual orientations flanking (i.e., ± 45°) the stimulated orientation on each trial, which compose the most likely incorrect interpretations of the evoked activity. For this, we projected single trial responses onto the decoder axes corresponding to the two orientations that were 45° apart from the presented visual input (see Methods). We found that SSTs suppressed evidence for flanking orientations exclusively in visually aligned trials (Figure 5F), while preserving correct (co-tuned) evidence. Thus, these results indicate that SST activation improves discriminability not by modulating population-averaged visual responses, but by decorrelating population-level representations via a specific reduction of evidence for flanking (competing) visual inputs.

Finally, if explaining away operates by reducing ambiguous activity, we reasoned that it should preferentially suppress co-tuned PCs whose activity (effects) also contributes to decoder evidence for competing orientations (causes). This latter group may include broadly tuned neurons, neurons tuned to intermediate orientations that fall between the stimulus and the second-best decoder prediction (similar to our model predictions), as well as neurons with more complex tuning functions^53^. To test this, we trained decoders using exclusively either co-tuned or anti-tuned PCs, relative to the input ensemble (Δθ(*SST*_*Ens*_ − *PC*) = 0° or Δθ(*SST*_*Ens*_ − *PC*) = 90°, respectively). Consistent with our predictions, co-tuned PCs showed a strong reduction in evidence for flanking (45° apart) orientations in SST-aligned trials (Figure 5G), while anti-tuned PCs were unaffected (Figure S4C). These results indicate that when SSTs, the visual stimulus, and the recorded PCs are all aligned, SST-mediated inhibition improves inference by selectively suppressing competing representations, consistent with a recurrent circuit-level implementation of explaining away.

Taken together, these results demonstrate that feature-specific inhibition aligned with matching visual inputs improves orientation coding primarily through competitive inhibition among similarly tuned PCs. Overall, our approach causally demonstrates how the functional microarchitecture of recurrent circuits augments the orientation-specific representational accuracy of the visual cortex.

## Discussion

In this study, we identify a recurrent excitatory–inhibitory–excitatory functional connectivity motif in mouse V1, mediated by orientation-tuned SST interneurons, that enhances the discriminability of visual inputs by precisely transforming afferent activity into more separable population codes. Using cell-type specific, all-optical circuit interrogation in awake mice, we functionally demonstrate that SSTs form a reciprocal inhibitory loop with PCs that is feature-selective. This PC→SST→PC motif enforces competition among similarly tuned PCs, explaining away partially redundant activity and improving the representational accuracy of co-tuned visual inputs.

Our findings help resolve longstanding debates over the role of recurrent circuits in sensory processing. Previous studies have yielded conflicting results as to whether photostimulating similarly tuned PC ensembles amplifies or suppresses local co-tuned representations^4,17,18,24,25,54^. By systematically varying input pattern size, we confirm that the same circuit can flexibly support both outcomes, with only very sparse ensembles eliciting net orientation-specific suppression and larger ensembles generally promoting completion. While feature suppression by ultra sparse activity might help enhance local differences in orientations in more complex natural scenes, our results also indicate that SST recruitment strongly and monotonically increased with the number of activated PCs, recapitulating previous one-photon experiments^44^. Indeed, we found that large PC ensembles maximally drove co-tuned SST activity. In turn, in the absence of excitation, co-tuned SSTs suppressed similarly tuned PCs. Because both inhibition and excitation converge onto similarly tuned PCs, the network outcome will flexibly depend on the precise recruitment these antagonistic forces^55^, as well as the network state^52^. Another advantage of this feature-specific negative feedback loop through PC→SST→PC connections is that it can minimize within-feature redundancy while preserving parallel, orthogonal codes.

Our results further clarify the distinct roles of PV and SST interneurons in shaping recurrent computations. Despite their very dense innervation by PCs^30,43^, PV neurons were suppressed by local PC inputs and exhibited no tuning-dependent responses. This motif may produce a spatial redistribution of inhibition in which SST activation suppresses dendritic activity in co-tuned PCs while indirectly relieving perisomatic inhibition via PV suppression. Such a circuit may selectively amplify “winning” inputs arriving to the soma, while suppressing predictable, statistically redundant components of the sensory drive in dendritic compartments^56^.

An alignment between excitation and inhibition of synaptic inputs in feature space has been observed across sensory systems^46,57–59^. Recent connectomics data further indicates that excitatory and inhibitory neurons converge onto common PC targets^60,61^, suggesting the existence of a canonical circuit motif. But what are the computational consequences of such opposing, feature-selective forces? We propose that this architecture allows different strategies of contextual modulation to optimize inference by either cancelling or completing predictable activity, depending on the structure of the input and the network state. While recurrent excitation among similarly tuned PCs has been proposed to amplify weak feedforward inputs and complete noisy or occluded representations^62–64^, unchecked recurrent excitation can increase redundancy, compromise discriminability^65^, and slow the processing of incoming inputs^55,66^. Here, we show that activation of SST ensembles sharpens the orientation tuning of matching visual inputs, improving their linear discriminability. Specifically, SST activation during visually aligned trials suppressed decoder evidence for the flanking visual orientations, consistent with competitive suppression of alternative interpretations. This pattern is a hallmark of “explaining away”, a computation in which networks enhance inference by actively suppressing partially overlapping, yet ultimately suboptimal, representations^11,12^. By selectively suppressing co-tuned PC activity that also contributes to competing interpretations, SSTs reduce representational overlap between similar stimuli and thereby enhance the linear separability at the population level. Like-to-like inhibition might thus promote perceptual phenomena like the tilt illusion^67^, in which differences in nearby visual orientations are exaggerated when presented in a center-surround configuration.

More generally, this circuit may represent a shared mechanism for sharpening cortical tuning in conditions where inputs contain redundant or predictable features, such as high-contrast, spatially extended and homogeneous inputs^10,13,68^. By enhancing competition among similarly tuned neurons, SSTs minimize redundancy and maximize discriminability within the local network. This motif may extend beyond sensory cortices: in higher-order areas such as the prefrontal cortex, a similarly structured inhibitory motif has been proposed to enhance choice-specific accuracy in a model of decision-making^65^. Thus, feature-specific recurrent inhibition may constitute a general computational strategy by which the cortex filters predictable information, suppresses ambiguity, and refines internal models of the world.

## Supplementary figure legends

**Figure S1.**
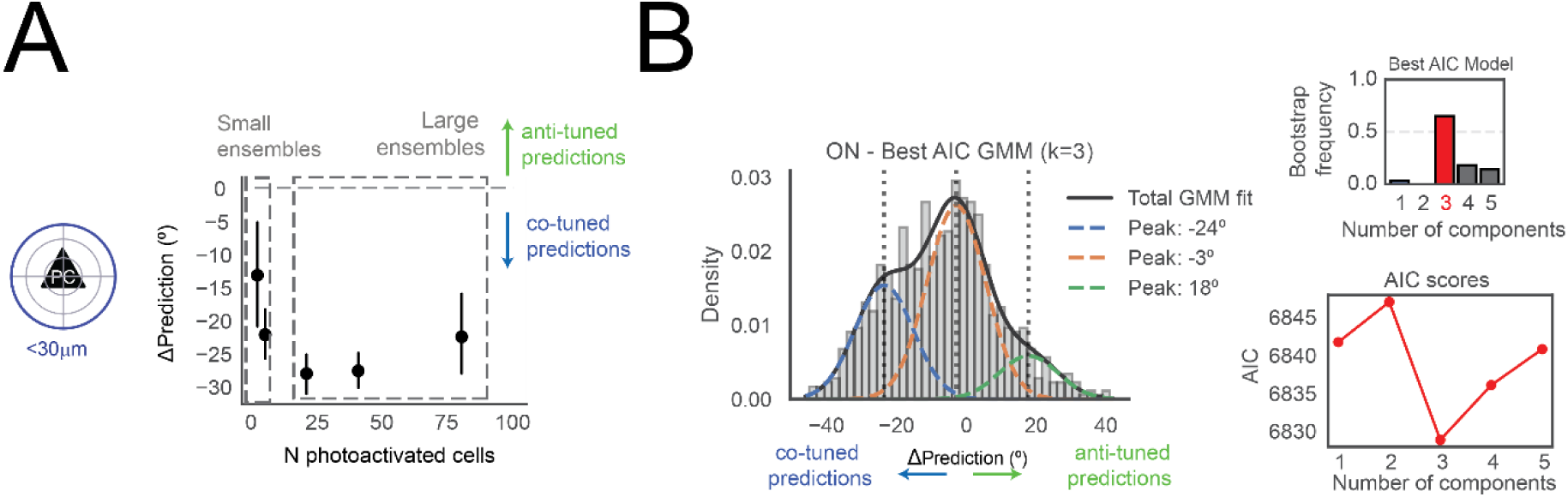
Decoder predictions from targeted and non-targeted neurons. **(A)** Mean ΔPrediction (°) computed using only the activity of cells within the photostimulated area, shown as a function of ensemble size. All ensembles yielded negative ΔPrediction values, indicating that even small groups of stimulated cells carried sufficient orientation-specific information to bias decoder output toward the target stimulus. Error bars represent 95% confidence intervals across ensembles. **(B)** Gaussian Mixture Model (GMM) analysis of ΔPrediction values from the non-targeted responses to all input ensembles. Top left: distribution of ΔPrediction and best-fit 3-component GMM with peaks at approximately –24°, –3°, and +18°, consistent with distinct network regimes (feature completion, neutral, and suppression). Bottom: model comparison using AIC shows that three components consistently provided the best fit. Top right: frequency with which each number of components was selected as the best model across 100 bootstrap replicates.

**Figure S2.**
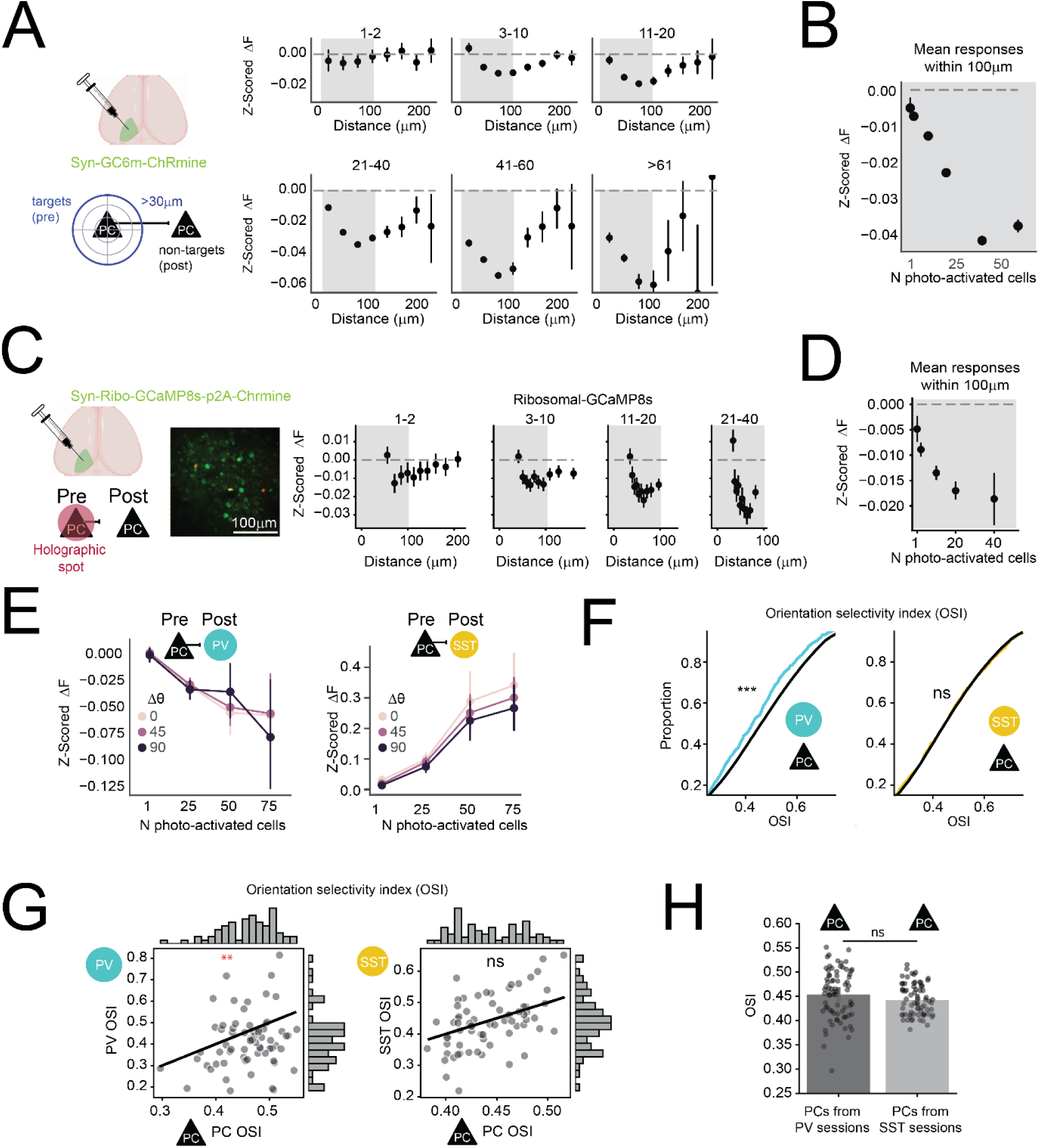
Distance-, size- and tuning-dependent responses of different cell types. **(A)** Left: Schematic of experimental design. Responses of non-target PCs (>30 µm away) were analyzed as a function of radial distance from the nearest target. Right: Mean z-scored and baseline-subtracted calcium responses (z-scored ΔF) of non-target PCs are plotted across increasing ensemble sizes, grouped by number of photostimulated cells (indicated above each panel). For all ensemble sizes, photostimulation produced distance-dependent suppression of nearby PCs, which gradually recovered at greater distances. Error bars represent 95% confidence intervals across ensembles. **(B)** Aggregated mean responses for distances within 100µm from the nearest photostimulated cell for each ensemble size. Error bars represent 95% confidence intervals across ensembles. **(C)** Left: Schematic and example image from a mouse expressing ribosomally localized GCaMP8s and Chrmine in PCs. Right: Mean z-scored and baseline-subtracted calcium responses (ΔF) of non-target PCs are plotted across increasing ensemble sizes, grouped by number of photostimulated cells (indicated above each panel). **(D)** Aggregated mean responses for distances within 100µm from the nearest photostimulated cell for each ensemble size. Error bars represent 95% confidence intervals across ensembles. **(E)** Joint distance- and tuning-dependent responses to PC ensembles in SSTs and PV interneurons. Left: In PV-Cre × Flex-tdTomato mice, photostimulation of PC ensembles suppressed PV interneurons. Right: In SST-Cre × Flex-tdTomato mice SST interneurons showed robust, size-dependent responses both for similarly tuned PCs (Δθ(*PC*_*Ens*_ − *SST*), = 0°, pink), and for anti-tuned PC ensembles (θ(*PC*_*Ens*_ − *SST*)= 90°, purple). **(F)** Cumulative distribution of orientation selectivity for PV, SST, and PC populations. PV cells show significantly lower orientation selectivity compared to surrounding PCs (***p < 0.001, Wilcoxon rank-sum test), while SST cells have similar selectivity to PCs (p>0.05). The x axis was cropped between 0.25 and 0.75 for visualization purposes. **(G)** Mean orientation selectivity for GABAergic neurons relative to local PCs. Each dot represents a session. Y-axis indicates the mean selectivity of either PVs (left) or SSTs (right). X-axis indicates mean selectivity of PCs. PVs, were reliably less orientation selective compared to nearby PCs (**p= 0.002, Wilcoxon test, N =75 sessions). SSTs were as orientation selective as nearby PCs (p>0.05, Wilcoxon test, N=69 sessions). **(H)** Mean orientation selectivity for PCs recorded from PV-tom or SST-tom mice. Each dot represents a session (p>0.05, Wilcoxon Rank Sum test, N>69 sessions).

**Figure S3.**
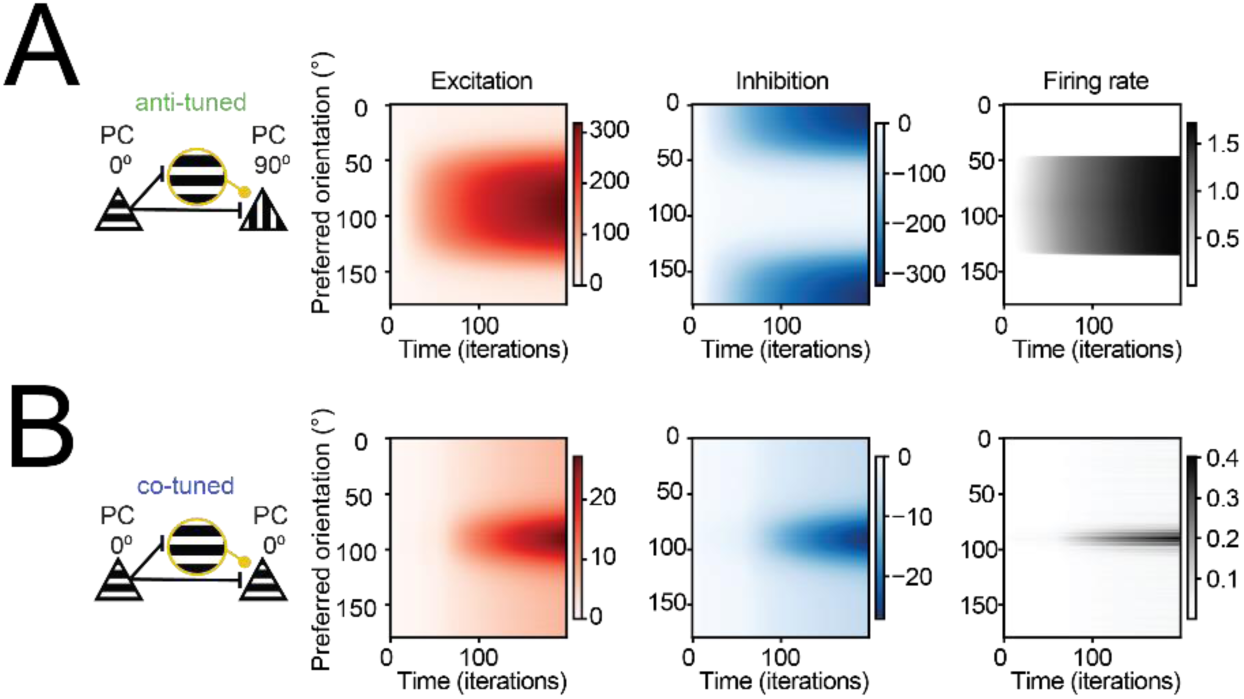
Temporal profiles of excitatory and inhibitory drives in models with opposite inhibitory connectivity. **(A)** Example simulation showing the evolution of excitation, inhibition, and firing rate for successive iterations within the recurrent loop in response to feedforward input, for a network with anti-tuned inhibition (top row, k_inh < 0). PCs are sorted by their preferred orientation. **(B)** Same as in **(A)** but for a model with co-tuned inhibition (k_inh > 0).

**Figure S4.**
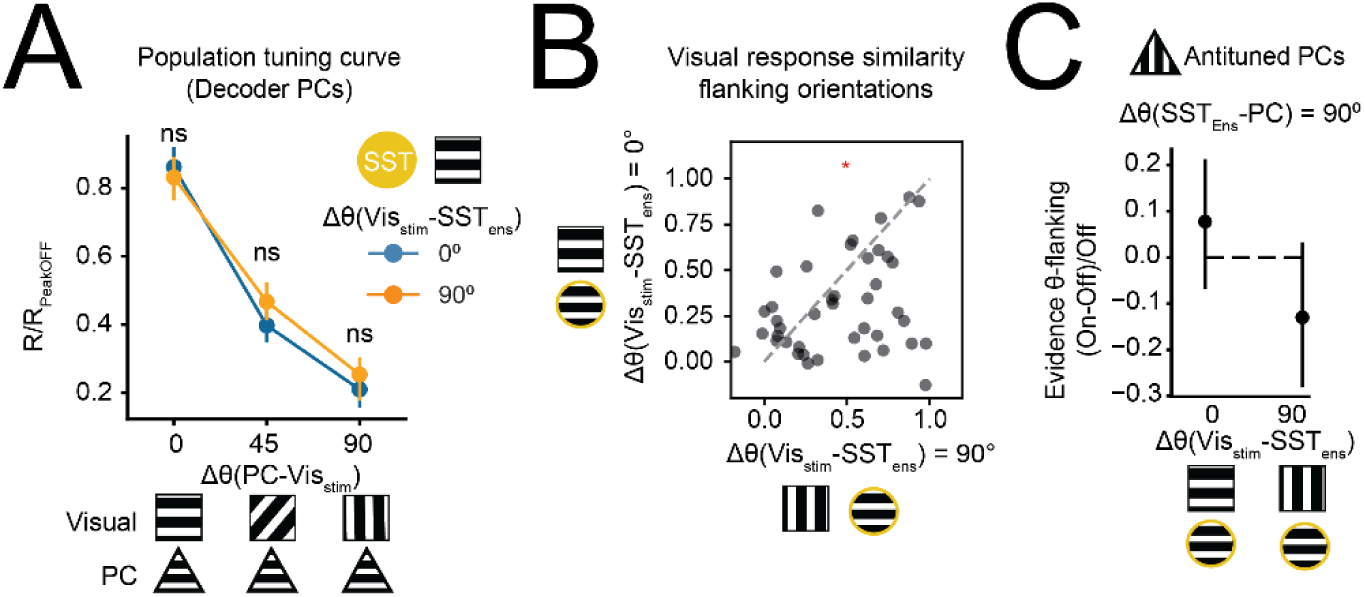
Visual response similarity to flanking orientations in co-tuned and anti-tuned PCs. **(A)** Population tuning curves of “decoder-informative” PCs (neurons with greater than zero L1 decoder weights) in SST aligned and SST misaligned trials. No differences were observed across any visual condition (p>0.05 for Δθ(*PC* − *Vis*_*Stim*_) 0°, 45°, or 90°). **(B)** Cosine similarity between responses to flanking visual orientations, normalized by the similarity between responses to the same visual orientation. Visually aligned SST ensembles enhance decorrelation of flanking visual responses relative to misaligned ensembles (*p=0.01, Wilcoxon test, N = 39 visual comparisons, 21 ensembles, 9 sessions, 6 mice). **(C)** SST activation does not modify the relative evidence for flanking orientations carried by anti-tuned PCs (Δθ(*SST*_*Ens*_ − *PC*)= 90°) (p>0.05, for both Δθ(*Vis*_*Stim*_ − *SST*_*Ens*_) = 0° and 90°, permutation test. N = 23 ensembles, 9 sessions, 6 mice).

## Methods

### Animals

All experiments were conducted in C57BL/6 mice of both sexes, aged 8 weeks or older. For experiments targeting pyramidal cells (PCs), we injected AAV8-hSyn-GCaMP6m-p2A-ChRmine-Kv2.1-WPRE into either PV-IRES-Cre;RCL-tdTomato, SST-IRES-Cre;RCL-tdTomato or EMX-IRES-Cre mice. Additional experiments were performed using AAV2YF-hSyn-ChRmine-p2A-riboGCaMP8s in PV-IRES-Cre;RCL-tdTomato mice (Figure S2). For experiments involving SST interneuron photostimulation, we used SST-IRES-Cre;RCL-tdTomato mice crossed with CaMKII-tTA;tetO-GCaMP6s mice and injected with a custom virus: AAV9-2YF-hSyn-DIO-GCaMP6m-p2A-ChRmine-Kv2.1-WPRE. In this configuration, SST interneurons were labeled with tdTomato via the Cre-dependent reporter and co-expressed GCaMP6m and stChRmine via the Cre-dependent viral vector. Concurrently, pyramidal cells expressed GCaMP6s via the tTA system. All procedures involving animals were approved by the Animal Care and Use Committee at the University of California, Berkeley.

### Surgery

AAV vectors were injected intracortically in V1 and cranial window surgeries were performed immediately after. Briefly, mice were anesthetized with isoflurane (2%) and administered 2 mg/kg of dexamethasone as an anti-inflammatory and 0.05 mg/kg buprenorphine as an analgesic. The scalp was removed, the fascia retracted, and the skull lightly etched. Following application of Vetbond (3M) to the skull surface, a custom stainless steel headplate was fixed to the skull with two dental cements: Metabond (C&B) followed by Ortho Jet (Lang). After the dental cement dried, a 3-mm diameter craniotomy over the left primary visual cortex was drilled, and residual bleeding stopped with repeated wet–dry cycles using sterile physiological solution, gauze, and Gelfoam (Pfizer). A window plug consisting of two 3-mm diameter coverslips glued to the bottom of a single 5-mm diameter coverslip (using Norland Optical Adhesive #71) was placed over the craniotomy and sealed permanently using metabond (Parkell). Animals were allowed to recover in a heated recovery cage before being returned to their home cage. Seven days after surgery, animals were habituated to head-fixation, and in vivo all-optical experiments were performed after at least 2 habituation sessions.

### Two-Photon All-Optical Read-Write Experiments

For all-optical read-write experiments presented in all fi gures scanless holographic optogenetics with temporal focusing (3D-SHOT) path was implemented in a Sutter MOM (Movable Objective Microscope, Sutter Instrument Co.), as described previously^34,35,69^. Ti:sapphire laser (Chameleon Ultra II, Coherent) was used for two-photon imaging, and femtosecond fiber laser (Satsuma HP2, 1030 nm, 2MHz, 350 fs, Amplitude Systems) was used for two-photon holographic stimulation. The holography path included a blazed diffraction grating for temporal focusing (600l/mm, 1000nm blaze, Edmund Optics 49-570), and a rotating diffuser to randomize the phase pattern and expand the beam. Both the imaging path and the holography path had spatial light modulators (SLM; HSP1920, 1920 X 1152 pixels, Meadowlark Optics). The imaging path SLM enabled optical axial focusing. The SLM was conjugated to the X resonant galvo and optical axial focusing was achieved by loading Fresnel-lens-like phase patterns on the SLM for each desired axial depth. The high refresh rate provided by the overdrive SLM allowed the acquisition speed to only be limited by the number of planes. Collected images were then cropped to remove the first 20 scanned lines to account for the phase transition time of the SLM. Imaging planes were spaced 27-35um axially and the number of recorded planes was 4-7 at 7.8-4.2 Hz respectively. The holography path SLM was used to display the holographic phase mask, calculated using the Gerchberg-Saxton algorithm to place several cell-sized diffraction-limited spots in 3D target positions. A zero-order block was placed in the holography path. The imaging path and the holography path were merged by a polarizing beam splitter before the microscope tube lens and the objective (Olympus 20X 1.0NA). To limit imaging artefacts introduced by the femtosecond laser, and from the visual stimulus, both the laser and the LED screen were synchronized to the scan phase of the galvo-resonant scanner (8 kHz) using two independent Arduino Mega (Arduino), gated to be only on the edges of every line scan. Precise alignment between imaging path and the holography path is achieved by a calibration procedure previously described in detail^69^. The match between the intended coordinates and the real targeted locations was quantified and corrected regularly by photobleaching specific 3D locations on a thinly coated fluorescent slide.

All-optical read-write experiments were conducted in 3 stages. First, we imaged visual responses of neurons in the V1 L2/3 FOV. Second, we performed an online analysis based on Suite2p to identify neurons responding selectively to each visual stimulus. Finally, we holographically stimulated these functionally identified neuron groups. For fast online analysis, each imaging plane was Suite2p’ed independently in parallel (‘multiprocessing’ package in python). On an 8-core CPU (Intel Core i7 10700K 3.8GHz) with 128GB RAM, online Suite2p of a 25 min imaging data (512 X 512 pixels) took 8 min. Then, all ROIs identified by the Suite2p were analyzed to identify functional groups of neurons. The top 100 most selective neurons (see below selectivity estimation) for each orientation were chosen for stimulation, from which groups of different total cells were made. Finally, the locations of the selected ROIs were used as the target coordinates by converting to SLM coordinates per the calibration described above. Each neuron was stimulated with a train of 10 to 20 pulses, 10ms long each, at a frequency of 10Hz. The total power under the objective for each cell was 7mW, which is the same power used for the physiological point spread function estimation.

### Visual Stimuli

Adafruit Qualia 9.7’ DisplayPort Monitor was placed 8 cm from the right eye of the mouse (2048 X 1536 pixels, 19.7 X 14.8 cm width and height). The backlight of the Adafruit monitor was triggered by the galvo-resonant scanner such that the monitor would emit light only during the mirror turnaround time, similar to the gating of the holographic stimulation described above. In addition, in all 2p experiments, electrical tape was applied between the objective and the mouse’s headplate to prevent monitor light from contaminating 2p imaging. For visual tuning blocks (no holography) oriented gratings of 4 orientations were presented using custom Bonsai rx software adapted from the International Brain Laboratory (IBL) pipeline^70^. Gratings parameters were: 80×60 visual degrees, 0.08 cycles/degree of spatial frequency, 100% contrast, 0-2 cycles/second of temporal frequency. Specifically, to avoid possible adaptation effects, SST photostimulation experiments during visual processing were combined with drifting gratings at 2 cycles/second. Both directions corresponding to the same orientation were taken as equivalent for target selection and posterior analysis. For static gratings, the spatial phase of the stimulus was randomly changed on each trial.

### Data analysis

Motion correction and calcium source extraction were performed using Suite2p ^71^. Briefly, raw calcium videos were motion-corrected with subpixel alignment = 10, and calcium sources were extracted. Calcium sources were accepted or rejected based on morphological and functional features defined per session, with manual examination to define thresholds and to identify red cells. Neuropil subtracted fluorescence vectors (F) (using a neuropil coefficient of 0.7) were used for downstream analysis. Calcium signals were acquired continuously, and each cell’s fluorescence was z-scored for each experimental block (either visual or holographic). Responses were then baseline subtracted on each trial. A typical trial lasted 3.5-5 seconds, with baseline periods of at least 1 second. Holographic targets were aligned to calcium sources by calculating the Euclidean distance between the centroids of all holographic targets and all calcium sources and finding the minimum. Rarely, the automated software assigned targets to calcium sources with distance >12 um; these were excluded from subsequent analysis.

### Orientation tuning estimation

To estimate the orientation preference and orientation selectivity of individual neurons, we analyzed their mean responses to four grating orientations (0°, 45°, 90°, and 135°). For each neuron, we computed the preferred orientation as the angle of the vector sum of responses in orientation space (180°).

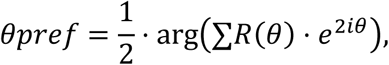

The orientation selectivity index (OSI) was defined as the magnitude of the normalized vector sum:

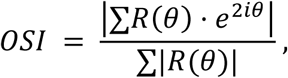

where *R(θ)* is the mean response at orientation *θ*, and θ is in radians. This metric ranges from 0 (untuned) to 1 (perfectly selective). For ensembles, we applied the same calculation to the mean response across all significantly photactivated ensemble neurons. Only neurons with OSI > 0.4 and ensembles with OSI > 0.3 were considered for tuning-dependent analyses.

### Decoder-based classification of visual-like activity

To evaluate whether holographically evoked patterns resembled visually evoked representations, we implemented a decoding-based assay. A linear support vector classifier (SVC; sklearn.svm.SVC, with “L1” penalty) was trained using baseline-subtracted population activity from non-targeted pyramidal cells recorded during visual stimulus trials. All non-targeted cells were included, regardless of their tuning properties. Four grating orientations (0°, 45°, 90°, 135°) were used for training. After training, the decoder was tested on holographically evoked trials in which no visual stimulus was presented. These trials involved optogenetic stimulation of ensembles of pyramidal cells tuned to a specific orientation. To quantify the alignment between the decoder’s output and the stimulated ensemble’s tuning, we computed the circular angular difference between the predicted orientation and the ensemble’s estimated preference. Since, for four equally spaced classes, theoretical chance-level prediction corresponds to a 45° error, we subtracted 45° from all ΔPrediction values to obtain a zero-centered chance performance (ΔPrediction ≈ 0°), with positive values (ΔPrediction > 0°) reflecting above-chance suppression and negative values (ΔPrediction < 0°) to reflect above-chance completion of the input orientation. To directly quantify the empirical Δprediction in the absence of any input for each recorded session, we decoded orientation from baseline periods (OFF) in each trial. These predictions served as controls for session-specific overrepresentation of orientations. We then compared decoder predictions on photostimulated (ON) versus OFF trials using paired scatterplots. Decoder outputs were compared across ensemble sizes using Wilcoxon signed-rank tests for paired differences in ΔPrediction. To assess whether the ΔPrediction distribution exhibited multiple underlying modes, we fit Gaussian Mixture Models (GMMs) with varying numbers of components (1–5) to bootstrapped samples of ensemble-evoked decoder outputs. Model quality was evaluated using Akaike Information Criterion (AIC), consistently identifying a 3-component GMM as the best fit across bootstrap replicates.

### Population-level decoder evidence

To quantify how holographic stimulation modulated the structure of orientation representations in the local network, we projected population activity onto the weight vectors (population axes) corresponding to each stimulus class of a linear decoder trained on visual-only trials (similar to ^17,24^). To evaluate the effect of holographic stimulation in figure 2, we computed the dot product between the trial-averaged activity vector evoked during photostimulation (no visual input) and each orientation-specific decoder weight vector. We summed this projected activity into a scalar “evidence” value for each of the four orientations, interpreted as the degree to which the evoked population activity aligned with the learned representations of each visual stimulus. We then computed the evidence as a function of the relative orientation between the visual stimulus and the stimulated ensemble. These evidence values were used to assess shifts in the distribution of stimulus representations under different ensemble sizes and tuning conditions.

### Rate-based Network Simulations

We implemented a simple recurrent rate-based network model comprising 180 neurons with uniformly distributed preferred orientations spanning 0° to 180°. Each neuron received a feedforward input generated by convolving a von Mises tuning curve with the stimulus orientation, using a concentration parameter (κ) of 10. To simulate natural variability, Gaussian noise was added to the feedforward input on each trial (standard deviation = 0.1).

Recurrent excitation and inhibition were defined by connectivity matrices computed from pairwise orientation similarity. The strength of excitatory-to-excitatory (E→E) and inhibitory-to-excitatory (I→E) connections was initially determined using orientation-tuned weight matrices based on the same von Mises function. We parameterized these matrices using independent concentration parameters for excitation (κ_exc = 4.0) and inhibition (κ_inh ∈ [–3.6, 3.6]). All diagonal elements in the weight matrices were set to zero, ensuring no self-connections.

To model the contribution of untuned background excitation and inhibition, we included additional random weight matrices (W_untuned_exc and W_untuned_inh) generated from a Poisson-distributed connectivity mask (λ = 3), scaled to ±0.1 and excluding self-connections.

Network activity evolved over 200 discrete time steps using a leaky update rule:

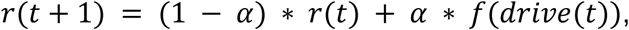

where α = 0.001 and *f(x)* is a bounded sigmoid activation function:

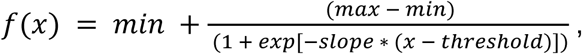

with min = 0, max = 10, slope = 3, and threshold = 2.

At each time step, the total synaptic drive to each neuron was computed as:

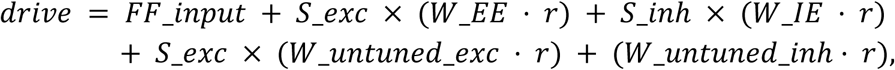

where S_exc and S_inh are dynamic scaling factors that modulate the gain of recurrent excitation and tuned inhibition, respectively, based on the current number of active neurons (i.e., neurons with activity > 0.01). These scaling factors were computed using a logistic function:

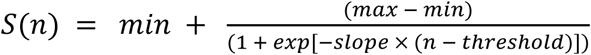

For excitation, we used parameters [slope = 0.3, threshold = 20, min = 0.05, max = 1.0], producing gradual growth with later saturation. For inhibition, we used [slope = 0.3, threshold = 20, min = 0.1, max = 1.0], generating rapid early recruitment and fast saturation.

Across each value of κ_inh, we simulated population responses to stimulus orientations (0–180°, in 5° steps). For each simulation, we recorded the feedforward input and the final network activity vector, and computed pairwise cosine similarity between stimulus conditions to assess how recurrent dynamics shaped output separability. We also estimated the half-width-half-max (HWHM) and the peak values for each output vector.

### SST Ensemble Stimulation During Visual Processing

To compare PC responses across conditions, we first computed z-scored and baseline-subtracted calcium traces for each cell and trial, as for all other analysis. Then, for each PC, visual responses in the unperturbed (SST-OFF) condition were examined across all stimulus orientations. If the minimum response was negative, an offset correction was applied by adding the absolute value of that minimum response, ensuring all values were non-negative. All responses were then divided by the peak response across orientations in the unperturbed condition. This normalization preserved the shape of the tuning curve while avoiding signs changes during division, enabling consistent comparison of relative tuning sharpness and suppression effects across stimulation conditions. Normalized responses were grouped by the angular difference between each PC’s preferred orientation and the presented visual stimulus, and averaged across experiments to generate population tuning curves (Figure 5C, Figure S4A).

### Decoder-based analysis for SST activation during visual processing

As in prior figures, a linear support vector classifier (SVC, with an “L1” penalty) was trained using population responses on an independent block of visual-only trials to predict stimulus orientation. This trained decoder was then applied to interleaved trials with or without SST ensemble stimulation, all with visual input. The neurons used to train and test the decoder were in all cases non-targeted PCs outside the photostimulation zone. All non-targeted cells were included, regardless of their tuning properties. Decoder error was calculated as the angular distance between the predicted and the true stimulus orientation. For each SST ensemble, we computed the percentage of change in decoder error between OFF and ON conditions, and correlated it with Δθ(*Vis*_*Stim*_ − *SST*_*Ens*_). Decoder performance was evaluated as a function of both Δθ(*Vis*_*Stim*_ − *SST*_*Ens*_) and statistically compared across SST-aligned and misaligned conditions. Correlation analyses were performed using Spearman correlation coefficient.

Decoder evidence for each orientation was extracted by projecting single-trial activity onto decoder axes. The “explaining away index” was defined as the reduction in evidence for the orientations flanking the visual stimulus presented on each trial (relative to visual-only trials) and used to assess suppression of competing interpretations.

To assess whether decoder-informative population vectors were decorrelated in different photactivation conditions in Figure S4B, we compared responses to each visual stimulus under SST photoactivation (ON) with responses of the same neurons to flanking visual stimuli without SST photoactivation (OFF). These responses were normalized by the similarity between matching conditions (same orientation, ON vs OFF visual responses). This ratio was then compared between SST-aligned and SST-misaligned visual inputs.

To identify the neuronal subpopulation mediating this suppression, independent SCV decoders were trained to each subset of co-tuned or anti-tuned PCs and the analysis was repeated for each of these subgroups.

### Statistical Testing

For group comparisons involving continuous variables (e.g., activity across tuning bins), normality was first assessed using the Shapiro-Wilk test. If any group failed to meet normality criteria (p < 0.05), we applied a Kruskal–Wallis H-test for multi-group comparisons or Mann–Whitney U tests for pairwise tests. For multiple group comparisons, we additionally performed FDR (Benjamini-Hochberg) correction. Additionally, for key comparisons we used permutation tests with 10,000 random shuffles to derive empirical null distributions. All plotted aggregated data represents the mean ± 95CI estimated via bootstrapping, unless otherwise noted. **Permutation tests:** To test for significant group differences without assuming parametric distributions, we implemented a two-sided permutation test. Trial labels were then shuffled 10,000 times to generate a null distribution, and compared mean or median differences. **Correlation analyses:** To assess trends between continuous variables (e.g., activity vs. relative orientation), Spearman’s rank correlation was used. **Paired comparisons:** To compare matched ON versus OFF conditions (e.g., SST ON vs. OFF for the same ensemble), we used paired statistical tests (either Wilcoxon signed-rank test or a paired Student’s t-test) depending on data distribution and assumptions.

## Acknowledgments

This work was supported by National Institutes of Health (NIH) grant R01-EY023756 (HA), U19NS107613 (HA), RF1-NS128772. H.A. is supported by a Chan-Zuckerberg Biohub Investigator Fellowship. M.B.O. was supported by the Weill Neurohub postdoctoral fellowship. This content is solely the responsibility of the authors and does not necessarily represent the views of the National Institutes of Health. We thank Janine Beyer for technical support. We thank Mei Li, supported by the NIH core grant EY003176, for custom viral preparations. We thank Lucia Rodriguez, Daniel Feldman and Antonia Marin-Burgin for their thoughtful comments on the manuscript. We also thank Kenneth Miller, Ho Yin Chau, and Agostina Palmigiano for valuable discussions on the data and computational analyses.

## Notes

### Competing Interest Statement

The authors have declared no competing interest.

